# Splice-site Strength Estimation: A simple yet powerful approach to analyse RNA splicing

**DOI:** 10.1101/2020.02.12.946756

**Authors:** Craig Dent, Shilpi Singh, Shikhar Mishra, Nawar Shamaya, Kok Ping Loo, Rucha Dilip Sarwade, Paul Harrison, Sridevi Sureshkumar, David Powell, Sureshkumar Balasubramanian

## Abstract

RNA splicing, and variations in this process referred to as alternative splicing, are critical aspects of gene regulation in eukaryotes. From environmental responses in plants to being a primary link between genetic variation and disease in humans, splicing differences confer extensive phenotypic changes across diverse organisms^1–3^. Current approaches for analysing splicing rely on quantifying variant transcripts (i.e., isoforms) or splicing events (i.e., intron retention, exon skipping etc)^4, 5^. However, regulation of splicing occurs at the level of selection of individual splice sites, which results in variation in the abundance of isoforms and/or splicing events. Here, we present a simple approach to quantify the strength of individual splice sites, which determines their selection in a splicing reaction. Splice-site strength, as a quantitative phenotype, allows us to analyse splicing precisely in unprecedented ways. We demonstrate the power of this approach in defining the genomic determinants of the strength of individual splice-sites through GWAS. Our pilot-GWAS with more than thousand splice sites hints that *cis*-sequence divergence and competition between splice-sites and are among the primary determinants of variation in splicing among natural accessions of *Arabidopsis thaliana.* This approach allows deciphering the principles of splicing, which in turn has implications that range from agriculture to medicine.

## Introduction

During mRNA formation, certain sections of the transcribed RNA (introns) are removed with joining of the adjacent regions (exons) in a process known as splicing. Splicing is a fundamental process in eukaryotic gene regulation^6–8^. Introns removed during the splicing process typically harbour canonical sequences at the 5’ beginning (GT) and the 3’ end (AG), which facilitate their identification as splice sites^9–11^. However, non-canonical splice sites and alternative GT or AG sites could also be used for splicing^12^. Selection of splice sites is a critical determinant of the products of splicing. Differential selection of splice sites gives rise to Alternative Splicing (AS), conferring both proteome diversity and phenotypic plasticity^6, 13^. Changes in splicing can modulate phenotypes across diverse organisms including sex-determination in flies, stress response in plants and genetic diseases in humans^1, 3, 6, 13–15^.

Current approaches to analyse splicing involve exploration of the transcriptome with next-generation sequencing, still mostly with short-read RNA-seq data^4, 5, 16, 17^. These approaches rely on mapping the short-read sequences to the gene and subsequently take two distinct strategies. The first strategy relies on quantifying splice variants (isoforms) based on the distribution of the reads observed in the RNA-seq data (Fig. 1a). Accurate estimation of isoform abundance requires a comprehensive understanding of the potential transcript isoforms, which is often lacking. In fact, there are efforts to obtain comprehensive reference transcriptome, which could help in such analysis^18^. In addition, short reads give evidence only for a small fragment of a transcript, and many regions do not differ between isoforms. Therefore, it is impossible to accurately assign every read to its parent isoform^16, 19^. As a result of this, reads that may come from a novel isoform, could be wrongly attributed to a known isoform, which can lead to misleading biological interpretation^20, 21^.

**Figure 1.**
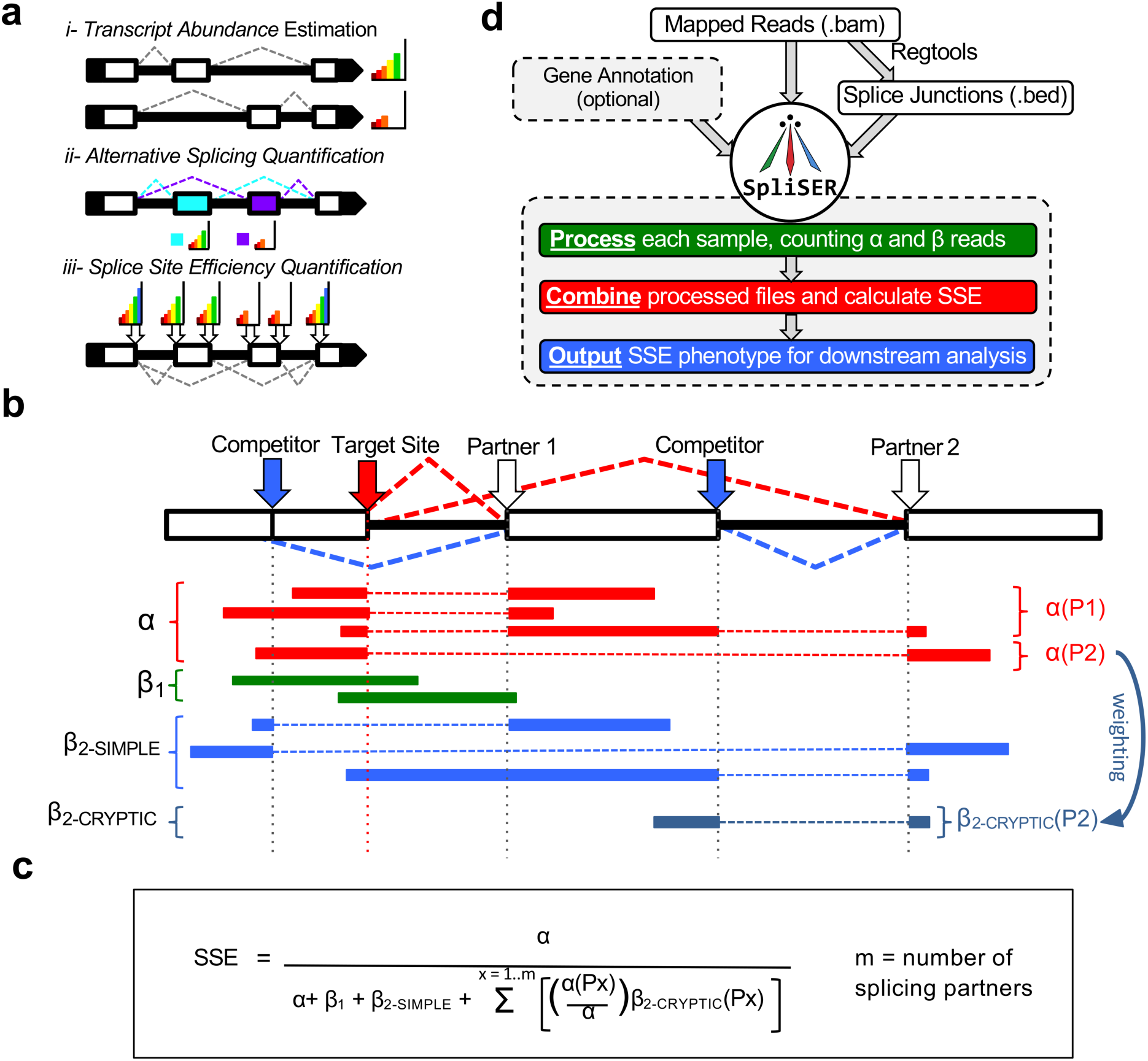
Overview of SpliSER and Splice-site Strength Estimation. a) Current methods for quantifying splicing from RNA-seq estimate isoform abundance (i), or compare the relative frequency of splicing events (ii). SpliSER quantifies the utilisation of individual splice sites (iii). b) For a given target splice site (large red arrow) Splice-site Strength Estimate (SSE) is calculated as the ratio of reads showing of reads showing utilization of the splice site (α-reads; red rectangles) and those showing non-utilisation (β_1_- and β_2_- reads; green and blue rectangles). α-reads (red) are split-reads with a gap in mapping which terminates at the target splice site, showing utilisation. The other splice site in this read is then considered a Partner of the target site. Two types of α-reads for a splice donor site with two different acceptor sites are shown. β_1_ reads (green) map either side of the target splice site but show no gap in mapping at this position, showing the target site has not been utilized. β_2_-reads (blue) show known Partners of the given site utilizing another splice site, which has therefore outcompeted the target site. β_2SIMPLE_ reads provide direct evidence of non-utilisation while β_2CRYPTIC_ reads need to be inferred. Thus, in calculating SSE β_2CRYPTIC_-read counts are weighted on the basis of the relative frequency of the partner through which they compete. c) Formula for arriving at SSE. d) Computational workflow of SpliSER. First, mapped reads are processed with Regtools to generate a bed file containing each detected splice junction. These two files are then provided as input to SpliSER, along with an optional annotation file defining gene boundaries.

An alternative approach evolved from a historical description of various types of splicing events such as intron retention, exon skipping, mutually exclusive exons and 5’ and 3’ alternative splice sites etc^22, 23^. Several approaches including rMATS^24^, ASD^25^, SUPPA2^26^ rely on quantifying splicing events. However, grouping splicing events across different genes to make inferences (e.g., increase in exon skipping or intron retention in a mutant compared to wild type) suggests biological categorization and common regulatory mechanism, which can be misleading. More recently, complexity-aware approaches such as MAJIQ^27^, Leafcutter^28^ and Whippet^4^ have taken alternative approaches, which somewhat minimise these impacts.

As described originally^22, 23^, the strength of the splice site defines patterns of splicing that governs the type of mRNAs produced. We reason that regulatory decisions on splicing occur at the level of individual splice sites rather than at the level of splicing events/isoforms. The extent of utilization or non-utilization of splice sites decides the type of transcripts produced. Splice-site selection affects individual (donor or acceptor) rather than pairs of splice sites (donor/acceptor), which leads to variation in splicing. For example, the same 5’ donor GT site can have alternative acceptor AG sites, which uncouples pairs of splice sites in their use. Therefore, there is a need to assess splicing at the level of individual splice sites if one has to understand the regulation of the specificity of splicing. Nevertheless, to date, there has not been much attempt to estimate the degree of utilization of any given splice site.

Here we present a conceptually different approach to analyse splicing and provide its bioinformatic implementation. We describe SpliSER (<u>Spli</u>ce-site <u>S</u>trength <u>E</u>stimate from <u>R</u>NA-seq), a bioinformatic tool to carry out *denovo* identification of individual splice sites from the RNA-seq data and derive a quantitative measure of their utilization. This approach moves away from analysing splicing at the level of isoforms or events. Splice-site Strength Estimates (SSE) increase the power to analyse regulation of splicing variation across the genome at a fine-scale resolution. We show an implementation of SpliSER to carry out GWAS analysis that allows us to detect SpliSE-QTLs, using the SSE as a quantitative phenotype. As a proof-of principle, using a pilot-dataset with 1500 sites that represent genes known to undergo alternative splicing and nonsense mediated mRNA decay^29, 30^, we map and show that with *cis*-regulatory variation and competition between splice-sites are among key determinants of differential splicing in these genes among natural accessions of *Arabidopsis thaliana*. Our strategy provides a powerful approach to decipher regulators of splicing and is widely applicable across diverse organisms.

## Results

We developed the idea of quantifying the usage of a given splice site earlier^21^, which we now refer to as Splice-site Strength Estimate (SSE). For any splice site (i.e., splice donor site), there are 4 types of reads that provide information about its usage (Fig. 1 a & 1b). First there are split reads that map perfectly with a gap between splice site and its partner sites. We call these reads as α-reads that provide evidence for the use of that splice site (Fig. 1b), independent of its partners (i.e., acceptor sites). Second, there are reads that cover the exon-intron junction (β_1_ reads), which provide evidence of non-usage of the target site (Fig.1b). We then use the historical concept of “competing splice sites”^31, 32^ to define additional reads that provide evidence of non-usage. If two donor sites partner with the same acceptor site, obviously both events cannot occur in the same transcript and thus the evidence for effective usage for one site becomes the evidence for the inefficient usage of the other site. These reads fall in two categories. Reads that define competitive splicing while also giving evidence for non-usage of the target site (β_2-SIMPLE_) or reads that define competitive splicing without read coverage of the target site (β_2-CRYPTIC_). While considering the total number of β_2_ reads, we apply weightings on the β_2-CRYPTIC_ reads. Thus, SSE is derived as the ratio of the evidence of utilization over the evidence of total possible utilization (Fig. 1c).

We further developed this concept into a bioinformatics pipeline, which we call as SpliSER (Fig. 1d). SpliSER uses the list of junctions to identify the splice sites from the RNA-Seq data. SpliSER, *per se*, is annotation-independent, yet allows the use of annotation when available. SpliSER has three different steps in its calculation of SSE. First, it uses a splice junction file to identify splice sites and records the counts of α reads for each of splice sites detected. Second, it provides the counts of β_1_ reads. Third, it analyses all the possible partners, quantifies them and provides a count of different β_2_ reads. Finally, it uses all of these metrics, applies filters (see detailed methods) and produces the SSE for all splice sites detected in the RNA-seq data.

To assess the accuracy of the SSE, we exploited natural variation in *Arabidopsis thaliana.* We reasoned that if our estimates are accurate, for accessions with mutations in splice sites, the SSE would be close to 0 for that site and will differ from other sites in the same gene. We identified accessions with splice-site mutations at *FLOWERING LOCUS M (FLM)*, which is known to undergo alternative splicing^33^ from the 1001 genome data. After confirming the mutations by Sanger sequencing, we downloaded the RNA-seq data for these accessions from the 1001 genome project^34^ and ran SpliSER across the *FLM* locus. Consistent with our predictions, we found the SSE of only mutated sites to be close to 0 unlike other sites known to be spliced with full efficiency (e.g., GT5, GT6, AG5, AG6, ^21^). In addition, the effect at a particular site was exclusive to accessions with mutations, confirming the specificity of our estimates (Fig. 2a). RT-PCR experiments confirmed alternative splicing, consistent with our estimates (Fig. 2b). Although total *FLM* expression level is substantially reduced in some of these accessions (Fig. 2b), we were able to estimate accurately, which indicates that we could pick up splicing differences independent of variation in gene expression.

**Figure 2.**
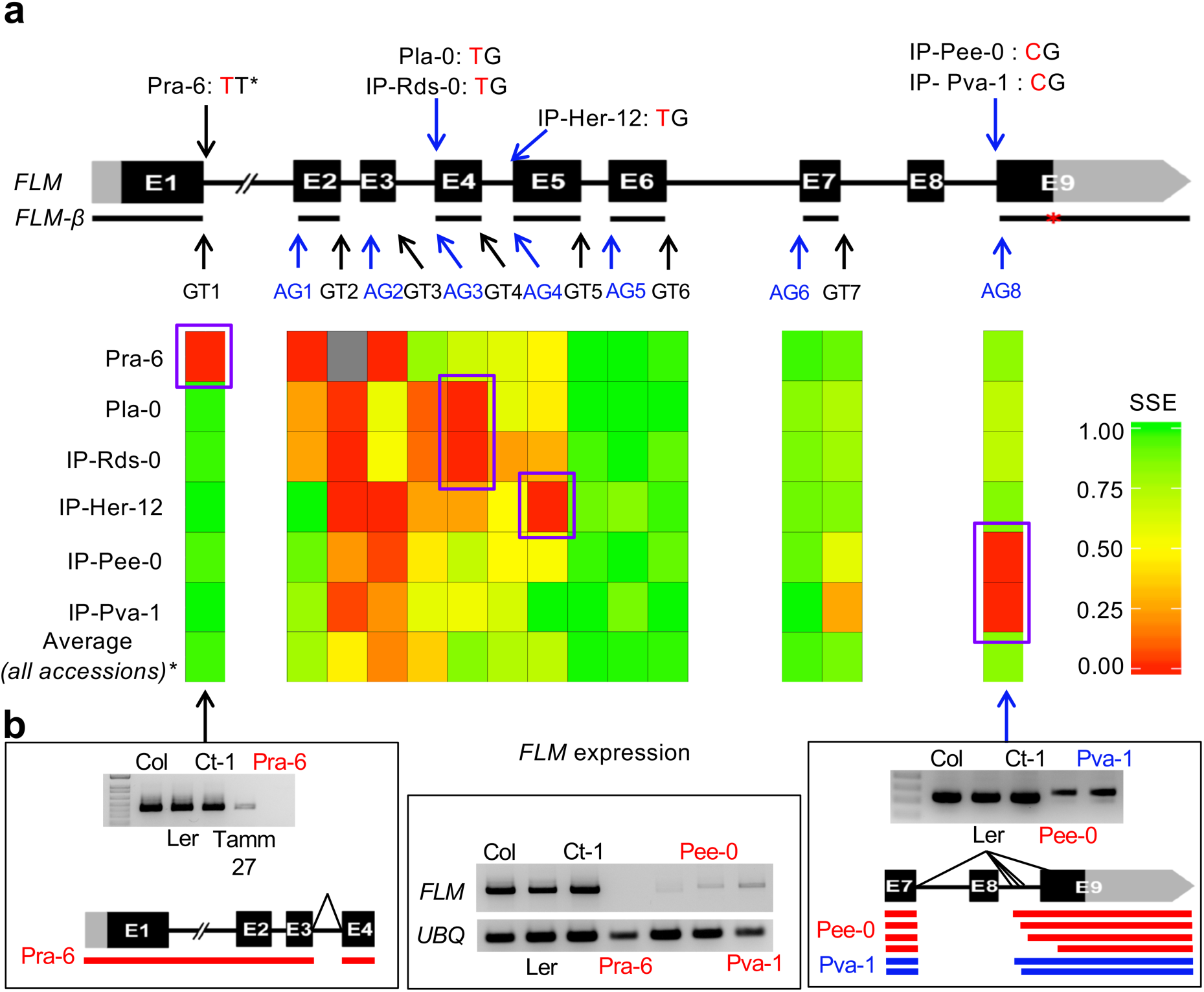
SpliSER estimates SSEs accurately. a) Gene model of *FLM* locus with splice site mutations across accessions. SSEs for the splice sites at the *FLM* locus in various genotypes. b) Experimental analysis with accessions harbouring splice site mutations at GT1 and AG8. The splice variants observed in the RNA-seq data are shown in the schematic. Splice site mutations at *FLM* are also associated with reduction in *FLM* expression.

SSE is a quantitative measure that can be used as a phenotype in GWAS to identify potential regulatory variation. To explore this possibility, we downloaded 6584 RNA-Seq datasets representing 728 accessions from the 1001 genome project^34^. Since we sought to compare the SSEs across accessions, we mapped all of these sequences against the TAIR10 Arabidopsis genome from the Col-0. The SpliSER pipeline was slightly modified for <u>SpliSE</u>-QTL analysis. SpliSE-QTL pipeline assesses whether there are at least 3 triplicate values for SSE for any given genotype/site, and then uses the BAMs that pass this filter to produce the SSE for a given genomic interval (e.g., genes or genomic position or chromosome) across all the accessions.

As a proof-of-principle pilot-study, here we carried out SpliSE-QTL analysis for all splice sites across 50 genes that are known to undergo alternative splicing and nonsense-mediated mRNA decay^29, 30^. SpliSE-QTL analysis detected close to 2000 splice sites across these genes. As expected, majority of the splice sites displayed minimal variation in SSE among accessions. However, there were sites with higher variability (Fig. 3a). To assess the genetic component of this variation, we calculated heritability of the SSEs. The heritability varied substantially (Fig. 3b), which indicated that there are splice sites with genetically controlled variability in SSEs. We focused on splice sites that were: a) in the upper quartile for heritability and variance and b) with minimum 100 accessions.

**Figure 3.**
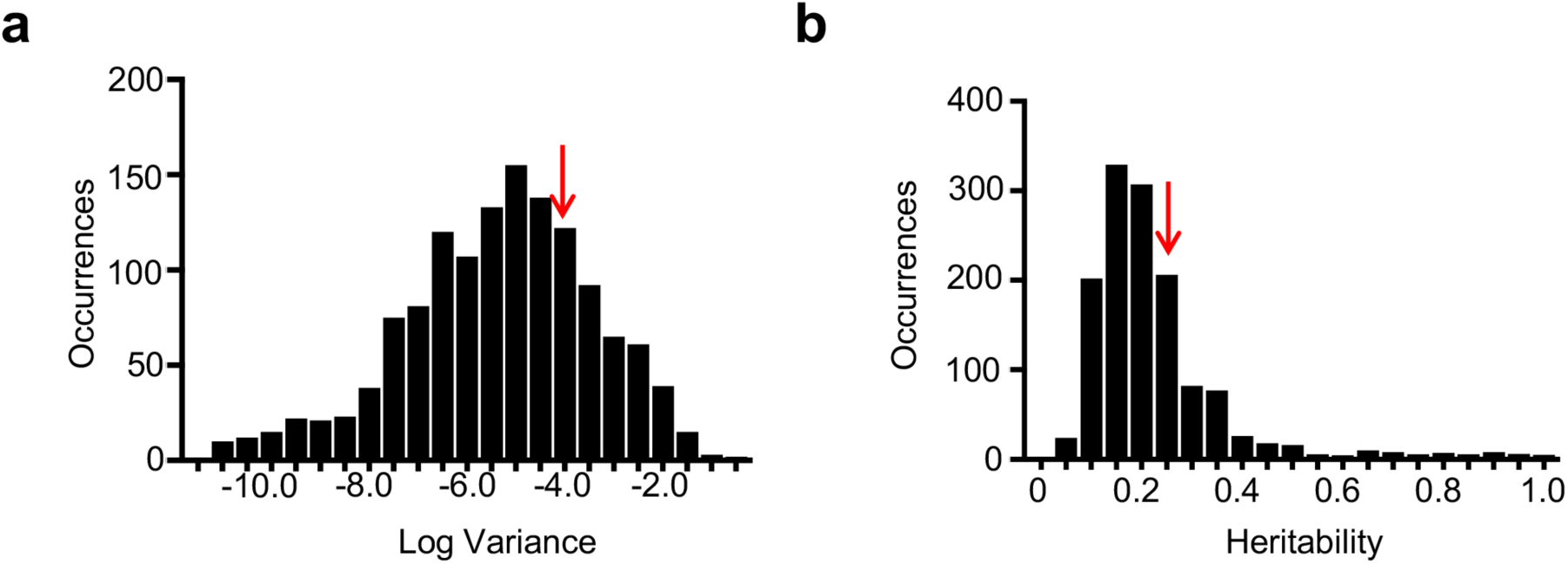
Splice-sites display heritable variation in their strength. a) Distribution of variances of the splice sites in AS-NMD genes. b) Distribution of heritability of the splice sites in AS- NMD genes. Sites to the right of the arrow represent in both a & b represent the upper quartile.

We identified significant, reliable associations for 46 sites that are spread over 19 of the 50 genes (∼38%) analysed in this pilot-study (Supplementary Table 1). To assess whether the GWAS signals are specific to splicing or simply reflected expression differences, we carried out eQTL analysis with the same set of accessions. We failed to detect overlapping GWAS signals for expression in most cases except for (*FLM (At1g770880)* & *AT4G357885),* potentially reflecting an association with NMD (Supplementary Table 2). To assess whether the SSEs are specific to splice-sites we compared the SSEs and GWASs with all SSEs in a single locus (*FLM*). SSEs differed between splice-sites within a gene, and not all sites gave a GWAS signal (Supplementary Table 3). Finally, similar associations were detected with competing sites (Supplementary Table 1). Taken together these findings suggest that the detected associations are specific to splice site and are associated with splicing differences rather than changes in gene expression.

To assess whether the associations hold up experimentally, we first analysed splicing of two sites at AT3G62190 that provided strong GWAS signals (Fig. 4a & 4b). SSE of two splice acceptor sites (canonical 23022226 & alternative 23022036) mapped to the same SNP (23022299). Common allele (G) at this SNP was associated with reduced strength of the canonical site and alternative allele (T) substantially increased the efficiency at the canonical site effectively competing out the alternative site (Fig 4b). Thus, accessions that differed at the associated SNP displayed distinct splicing patterns (Fig. 4c). We reasoned that if the observed associations are real, we should be able to predict the splicing patterns based on the genotype of the associated SNP. We tested this prediction using accessions that were not part of the GWAS panel. Consistent with our hypothesis, splicing patterns conformed with predictions (Fig. 4c). We carried out similar analysis with other loci and found them to conform predictions (Supplementary Fig S1-S3), which provided experimental support for detected associations.

**Figure 4.**
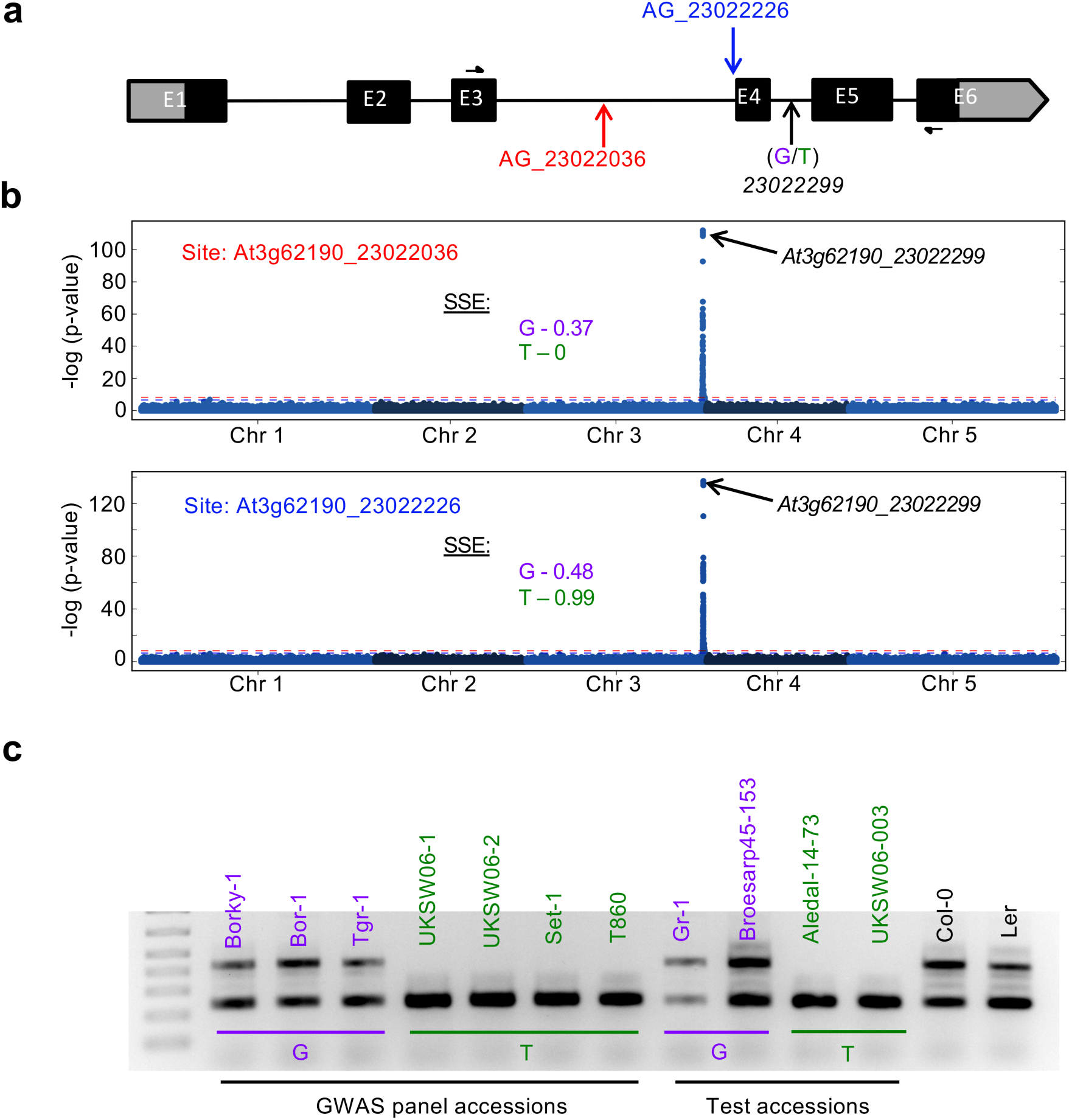
SpliSE-QTL analysis detects experimentally verifiable associations for splice-site strength. a) Gene model for At3G62190. Positions of two competing splice acceptor sites and the associated SNP are shown. b) GWAS analysis for SSE of both competing splice-sites identifies At3g62190_23022299 as the highest associated SNP in GWAS. The allelic effects on both sites are shown. c) Experimental verification of the SSE differences associated with 23022299-SNP. Common allele is shown in purple and the alternative allele is shown in green. Col and Ler harbour common alleles at this SNP.

Having confirmed the associations, we considered whether SpliSE-QTL has the ability to pick up causal mutations. To assess this, we analysed whether any of the detected associations actually mapped to the splice-site itself in this dataset. Variation in SSE for two sites (AT2G18300_7953316/17, and AT5G65050_25985653/54) mapped to the splice sites themselves (Fig. 5a). In addition, there were three more sites (AT5G46110_18699429/30, AT5G65050_25986012/13, AT5G65050/60_25989670/71), where the associated SNPs were within ±2bp of the splice site (Fig. 5a). While this is consistent with the idea that mutations at the splice site or in its vicinity would weaken the strength of a splice site, it also argues for the potential of SpliSE*-*QTL analysis to identify causal mutations underlying variation in splicing.

**Figure 5.**
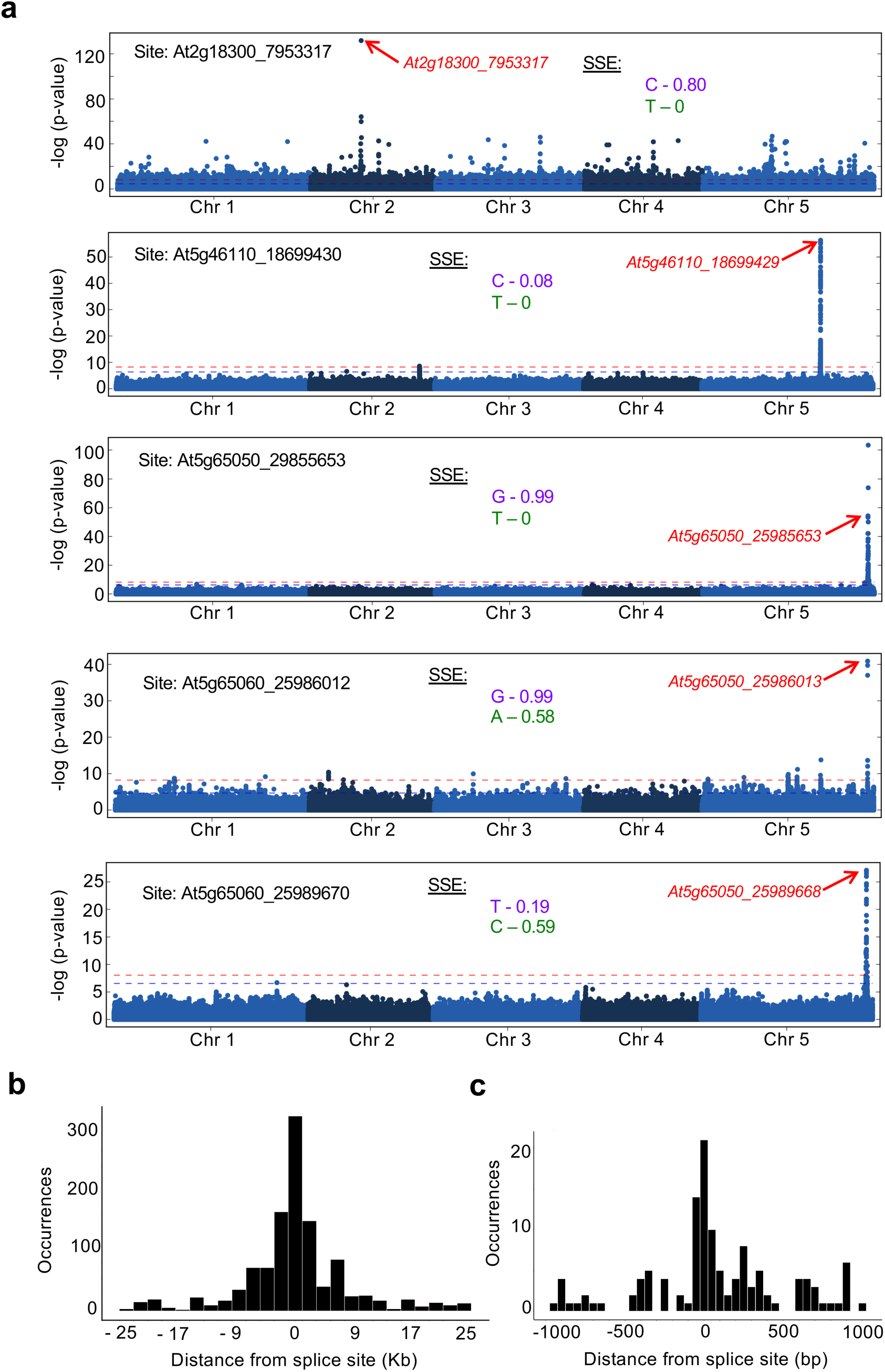
SpliSE-QTL can detect causal variants and *cis-*regulatory variation is among the primary determinants of variation in splicing. a) Manhattan plots for the splice sites in which the GWAS identified mutations in the splice-site itself (At2g18300_7953317, At5g65050_25985653) or in its vicinity (At5g46110_18699430, At5g65060_26986012 and At5g65060_25989670 harbouring alternative alleles at +/-2 positions, see Supplementary Fig S6- S11 for additional Manhattan Plots). The splice-site strength conferred by the common (purple) and the alternate (green) alleles are shown. b) Distribution of distances of Top 25 highest associated SNPs for the splice-sites for which associations were detected on the same chromosome. c) Distribution of the distances of the highest associated SNPs that fell within 1Kb region

Next, we analysed the pattern of detected associations. We noticed that most of the GWAS signals (44 out of 49) were on the same chromosome as that of the splice site (Supplementary Table 1). To assess the pattern, we compiled the distances of the associated SNPs (Top 25 SNPs for each phenotype) from the splice site (Supplementary Table 1, Fig. 5b). Distances clustered within 2Kb of the splice site (Fig. 5b). Most of the top-associated SNPs (32 out of 44) fell within 2Kb of the splice site (Fig 5c), suggestive of a strong *cis-*effect. In fact, for 26 out 44 associations, we detected an associated SNP within 100 bp from the splice site (Supplementary Table 1). We also noted several instances of common associated SNPs for multiple splice-sites, which suggested the influence of one splice-site over the other through competition. These findings indicate that in this dataset, *cis* rather than trans-sequence variation and competition between splice-sites are among the primary drivers of variation in splicing.

A recent study utilized the same RNA-Seq data from the 1001 genome project and carried out isoform-based splicing QTL analysis^35^ using sQTLseekeR^36^ based UlfasQTL^37^ analysis and concluded that majority of splicing variation in Arabidopsis is due to *trans* QTL^35–37^. This finding is in contrast to our current studies as well as earlier genetic studies^38^ that argued for *cis-*regulatory variation as primary determinants of splicing. Therefore, we compared the GWAS results obtained in this study with ours for the common genes (Supplementary Table 4). We observed that the isoform-based approach failed to detect even a single SNP that we report for the same set of genes (Supplementary Table 4). In addition, there was substantial noise in Khokhar et al’s GWAS study such that associations were found with SNPs scattered around the entire genome, while we detected specific signals (Supplementary Table 4, Supplementary Fig. S4). Thus, in addition to reducing noise, SpliSE*-*QTL analysis also provided specific associations with splice-sites.

Having confirmed the utility of SpliSER, we also implemented a method that would allow detection of differentially spliced sites across the genome between samples. To assess the efficacy of this approach, we applied SpliSER on a previously published RNA-Seq data on quintuple mutants of the *sc35-scl* in *Arabidopsis thaliana*^25^. In this comparison, Yan et al, utilised ASD to detect 213 differentially spliced events representing 199 genes (p-value<0.05) of which 34 were detected after FDR based analysis (FDR<0.05) ^25^. Applying SpliSER on the same data, we detected a total of 13094 splice-sites, representing 6457 genes to be differentially spliced (p-value<0.5), including 8 genes that were experimentally analysed by Yan et al (Supplementary Figure S5). Applying Benjamini-Hochberg correction, we detected 4431 sites representing 2688 genes. Of these 1380 sites representing 1012 genes displayed more than 20% change in splicing efficiency. Even this stringent list contained 28 of the 34 genes identified through FDR analysis by Yan et al (Supplementary Figure S5). Thus, diffSpliSE-analysis detected additional genes and provided splice-site based details of differential splicing.

To the best of our knowledge Splice*-*QTL is the only approach that allows detecting genetic variation that is associated with changes in splicing of a specific splice site. Our findings indicate that SpliSER provide reproducible quantification of SSE and SpliSE-QTL analysis has the potential to detect genomic determinants of variation in splicing. While our analysis is primarily based on short read RNA seq data, in theory the same principles could be applied with long-read data with minor changes. Our findings suggest that *cis-*regulatory changes and competition between splice-sites based on the splice-site strength could be among the key determinants of variation in splicing. While more work is needed to explain the mechanisms for each of these associations, we have demonstrated that these associated SNPs can be used as markers for tagging-splicing patterns. Splice-tagging SNPs would be of great use as markers having wide-ranging implications from agriculture to human disease. Future GWAS studies across genomes would unravel both the complexity and regulators of variation in splicing in an unprecedented fine scale.

## Online Methods

### Plant material/DNA/RNA Analyses

Seeds of the 1001 genome project accessions were obtained from European Arabidopsis Stock Centre. DNA and RNA extractions were done as described previously^39^. For gene expression studies DNAse I (Roche)-treated 1ug of total RNA was used for cDNA synthesis using the First strand cDNA synthesis kit (Roche) and the resulting cDNA was diluted and used for RT-PCR experiments. Primers used in RT- PCR analysis are given in Supplementary Table 5.

### Splice-site Strength Estimation

SpliSER requires a BAM file of mapped RNA-seq reads and BED file that contains a list of splice junctions detected in the alignment (such as those produced during mapping by TopHat2 or HISAT2, or directly from a BAM file using Regtools^40^). First, SpliSER uses the junctions BED file of each RNA-seq sample to define a list of splice sites observed in all samples; the read count of each junction is concurrently added to the α-read count for each of the two sites forming the junction. The list of “partners” (sites with which a given site has been observed to form a junction) is also recorded for each splice site. Second, SpliSER uses Samtools^41^ view command to retrieve reads whose mapping covers each nucleotide either side of the splice-site, the CIGAR string of these reads are then traversed to identify reads which map perfectly on either side of the splice site; these are counted as β_1_-reads for each site. Third, SpliSER traverses the list of partners of each splice site – any α-read belonging to the regular partner and rare partners, but not used in a junction with the splice site under consideration, is counted as a β_2-SIMPLE_ and β_2- CRYPTIC_ –reads respectively. During the calculation of SSE a unit interval weight is applied to the β_2-CRYPTIC_-read count coming from each partner site, representing the relative occurrence of that splicing partnership (e.g., If Site A is observed to partner with Site B 9 times, and Site C (whose regular partner is not Site A) once– the β_2-CRYPTIC_ reads from Site C will be assigned the weight 0.1. Thus Splice-site Strength Estimate is defined as

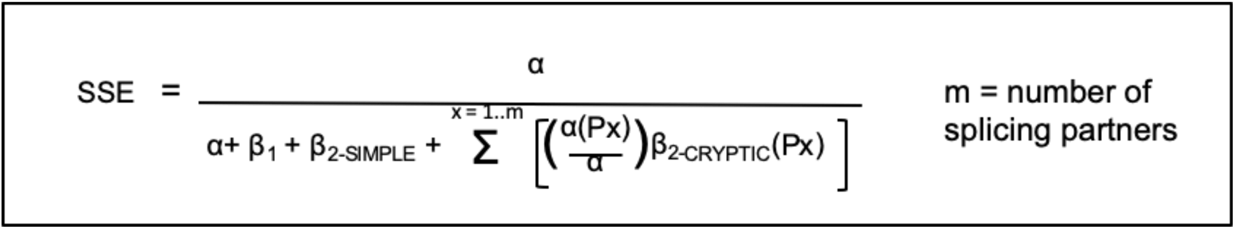

where β_2_^P^ is the β_2_ read counts coming from partner *P*, and *m* the number of partners. In each sample, for each splice site; SpliSER filters those with too few reads (sum of α, β1, and β2: Default 10). These parameters could be adjusted to increase the sample size and we found that decreasing the sum count to as low as 3 still provided similar signals in SpliSE-QTL analysis. **URLs:** *SpliSER* software, and scripts for statistical testing: https://github.com/CraigIDent/SpliSER

### SpliSE-QTL Analysis

For the SpliSE*-*QTL analysis, 6854 RNA-seq samples were downloaded from SRA, representing 728 accessions (PRJNA319904). Each sample was aligned to the TAIR10 reference genome using Tophat (v2.1.1; paramters --minIntronLength 20, --maxIntronLength 6000, -p 6)^42^. The resulting BAM files were indexed and sorted using Samtools sort v1.7^41^. The sorted BAM files and BED files for each sample were passed to SpliSER with a minimum of 10 evidencing reads required for a site to be called, and using a table adapted from the TAIR10_genes.gff3 annotation file to identify gene boundaries. The Splice-site Strength Estimate of each site in each of the 50 NMD target genes was quantified. The SSE phenotypes were further filtered using custom python scripts; an accession was only considered for further analysis for a given site if it had three or more individual RNA-seq samples underlying its average splicing efficiency value. A site was only considered for futher analysis if it contained 50 or more such accessions. We calculated the broad-sense heritability (H) and variance (v) of each splice site using a custom R script. GWAS experiments were performed using the easyGWAS^43^ and/or GWAPP^44^ web portal using the EMMAX/AMM algorithm with a minimum minor allele frequency of 5%, with no transformation of phenotypes. To assess the relationship between heritability/variance and the ability to detect GWAS signals, we initially carried out GWAS with all 1500 phenotypes. Our findings suggested that most of the signals were detected among splice-sites with a higher levels of heritable phenotypic variation. When variance is low, it resulted in spurious signals throughout the genome. Therefore, each of the Mannhattan plots were individually inspected, before deciding on whether a GWAS signal that shows statistical significance is trustable. In subsequent iterations, only sites that were in the upper quartile of heritability and variance were taken for analysis. In total more than 1500 phenotypes were analysed through GWAS that ultimately resulted in 49 associations for 19 of the 50 analysed genes (Fig.4, Fig. 5, Supplementary Figures S6-S11, Supplementary Table 1). All of these 49 were tested both through EMMAX algorithm or through AMM algorithm and both methods provided consistent signals. In instances where multiple SNPs that were in LD were giving same p-value of highest association, the closest SNP to the splice-site is presented in the Table.

### eQTL Analysis

eQTL analysis was carried out using published expression levels for the RNA-Seq data using the same panel of accessions that contributed to the splicing-QTL for each of the phenotypes, using the same GWAS options described above. The data were then summarised at the gene level and if any one of the panel of accessions gave a eQTL signal, it was considered to be a positive overlapping QTL. This analysis resulted in two genes (*FLM* and *At4g35875*) having overlapping eQTL and SpliSE-QTL signals.

### DiffSpliSE analysis

The six RNA-seq samples described in Yan et al^25^ were aligned to the TAIR10 reference genome using TopHat2 v2.0.10 (parameters -i 20 -I 6000 -g 10 -r 0 --mate-std-dev 50 --coverage-search, and indexed using Samtools v1.2. Each resulting BAM and junction BED file, were processed with SPliSER (command: process, parameters –m 6000) using an annotation file derived from the TAIR10 genes.gff3 file. The resulting SpliSER.bed files were then combined (command: combineDeep; parameters -1 Chr1) and output (command: output; parameters –t diffSPliSE, -r 10). Differentially spliced sites (DiffSpliSE) were detected with a custom R script. Briefly, we remove all sites containing an NA in any sample generated by insufficient read coverage; then filter all sites whose mean SSE (average of all samples) is <0.05 or >0.95, blind to experimental grouping. We utilized the EdgeR package^45^, testing for significant changes in splice site strength using a generalized linear model (glmLRT() function, default parameters) with a contrast corresponding to the difference between the α and β (β_1_ + β_2-SIMPLE_ + weighted β_2-CRYPTIC_) read counts, between two samples [ie. `(alpha.group1-beta.group1)-(alpha.group2-beta.group2)]`. Differentially Spliced sites were called as those with an FDR-corrected p-value <0.05, and an absolute change in averaged SSE >0.1 between conditions.

## Acknowledgments

We thank the European and North American Arabidopsis Stock Centres, and Alex Fournier-Level for the seeds. We thank Paul Harrison for his help and suggestions on statistics and Arun Konagurthu and Ya-Long Guo for advice on data processing and critical comments on the manuscript. This work is supported by the Australian Government’s Research Training Program (RTP) fellowship to CD, Australian Research Council –Future Fellowship (FT190100403) to SrS, ARC-Discovery Project DP190101479 and Chinese Academy of Sciences President’s International Fellowship for Visiting Scientists (CAS-PIFI 2019) to SB.

## Author contributions

CD, SrS and SB designed and conceived the study. CD, ShS, SM, KPL, PH, DP and SB developed the pipelines for SpliSER. CD, ShS, NS, RS and SrS carried out the experiments. SrS and SB acquired funding and SB conceptualized, coordinated and supervised the study.

## Data Availability Statement

All data used in this manuscript is publicly available and has been referenced appropriately in the manuscript.

## Code Availability Statement

SpliSER is available at GitHub at https://github.com/CraigIDent/SpliSER

## Supplementary Information

**Supplementary Figure S1.**
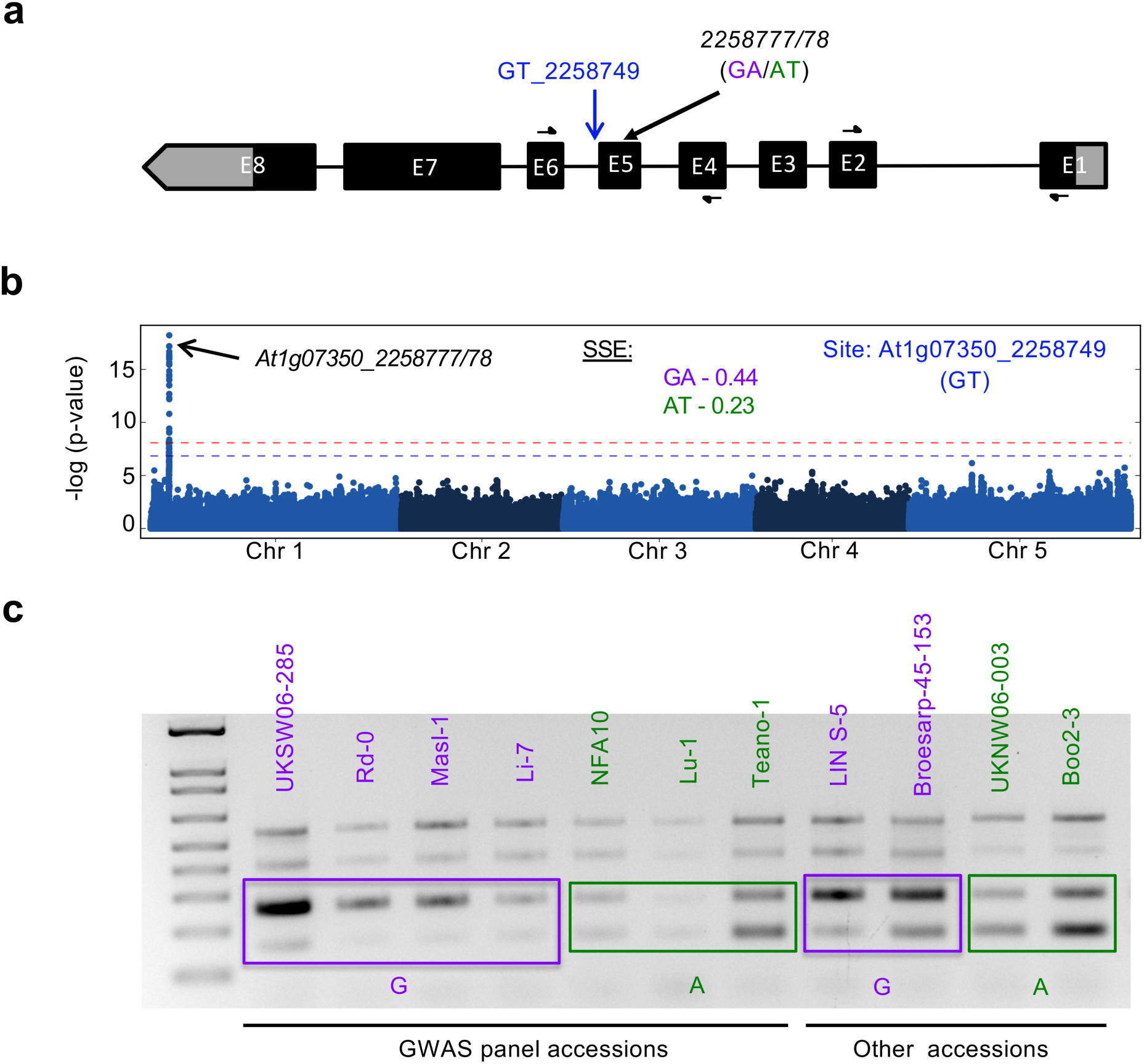
Experimental verification of associations for splice donor site At1g07350_2258749. a) Gene model for At1g07350. Positions of the splice-site and the position of the associated SNP. b) GWAS analysis for SSE of both splice-sites identifies At1g07350_225777 and 225778 (in LD) as highest associated SNPs in GWAS. The splice-site strength conferred by the common (purple) and the alternate (green) alleles are shown. c) The SSE differences associated with the associated genotype. Common allele is shown in purple and the alternative allele is shown in green.

**Supplementary Figure S2.**
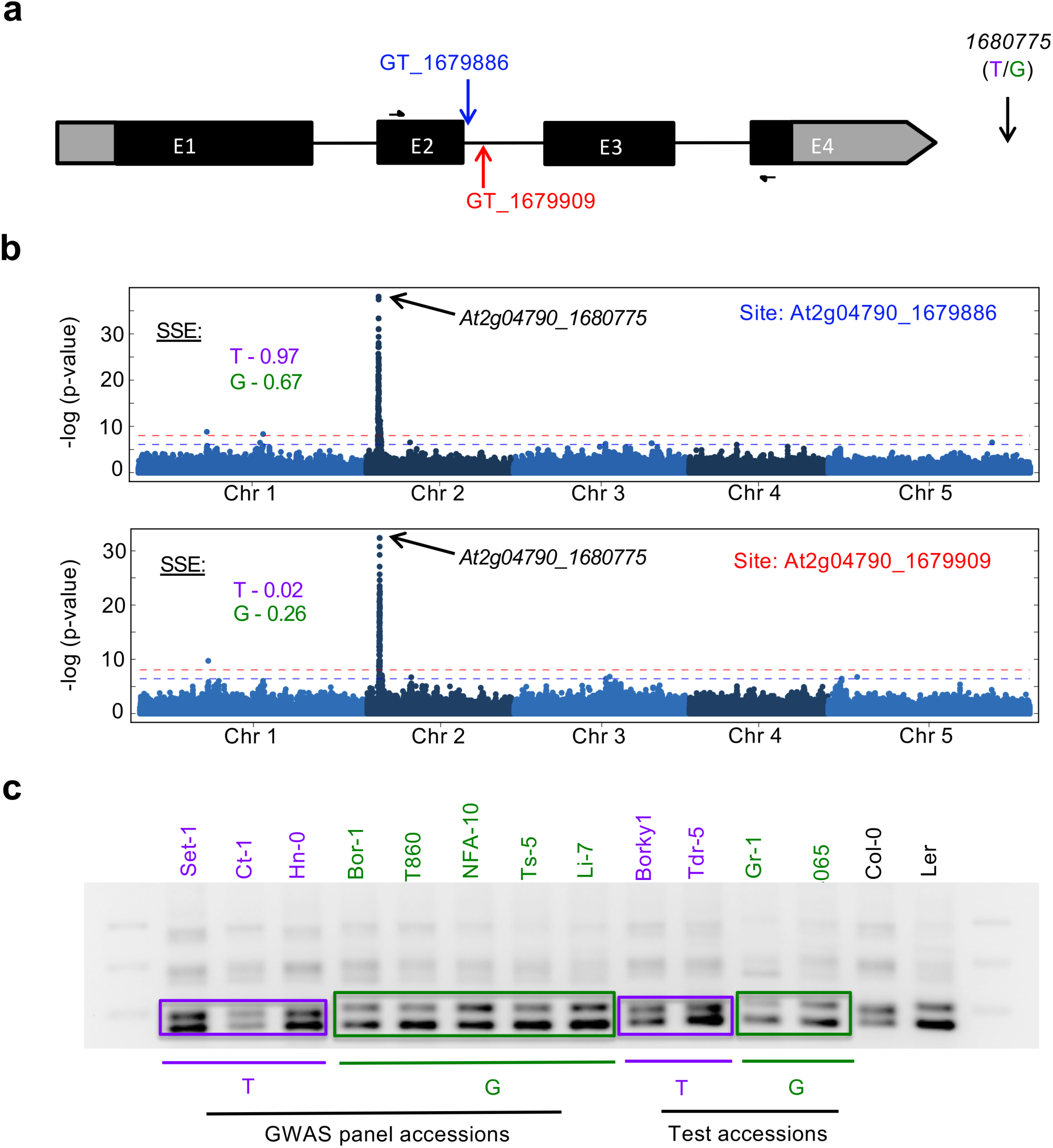
Experimental verification of associations for two splice-sites at At2g04790. a) Gene model for At2g04790. Positions of two competing splice-sites and the associated SNP (which is outside the gene boundary) are schematically represented. b) GWAS analysis for SSE of both splice-sites identifies At4g04790_1680775 as the highest associated SNPs in GWAS. The allelic effects on both sites are shown. c) The association of the SSE differences with corresponding genotypes can be experimentally verified. Common allele is shown in purple and the alternative allele is shown in green.

**Supplementary Figure S3.**
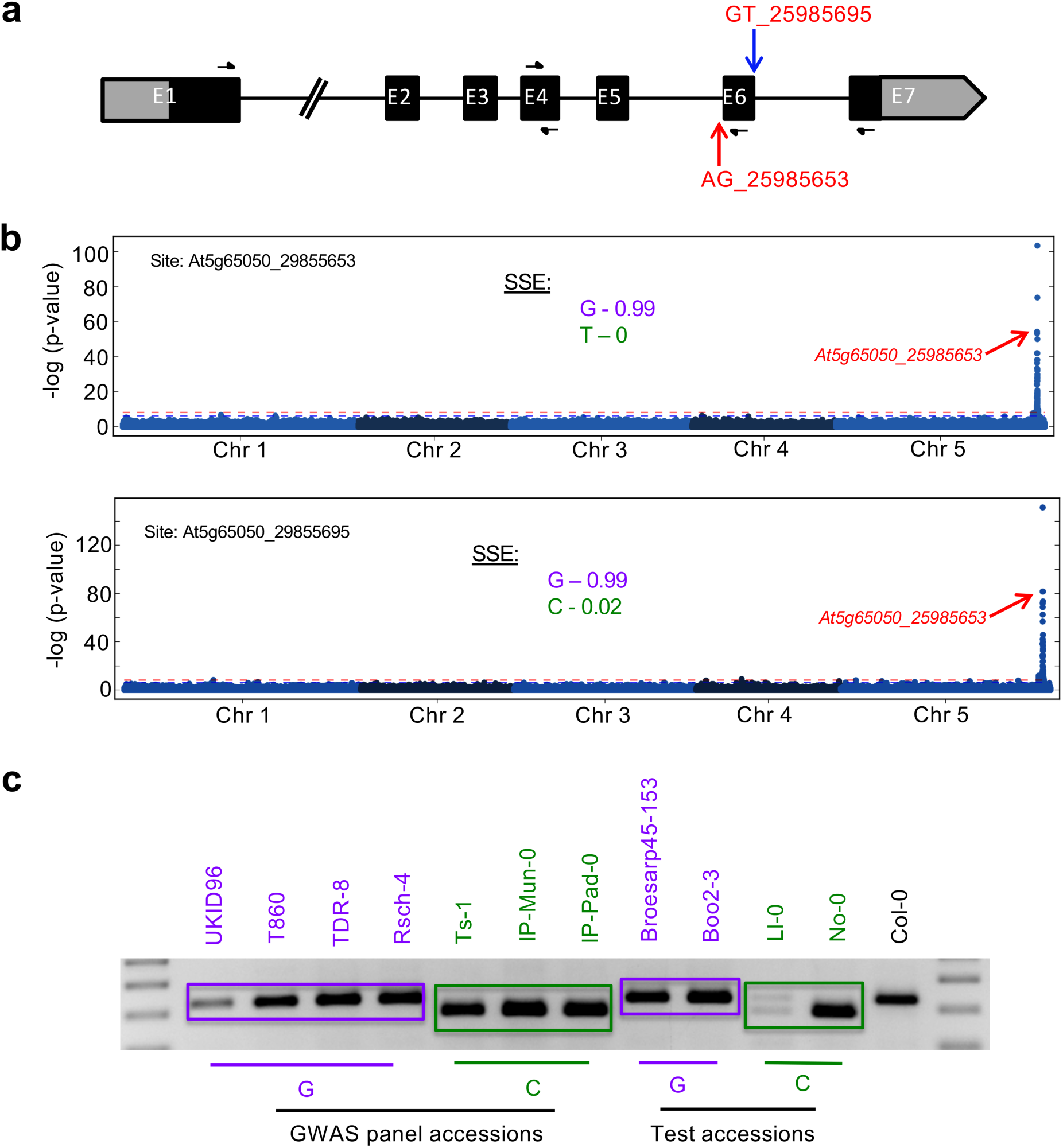
Experimental verification of associations for at At5g65050. a) Gene model for At5g65050 (MAF2). b) GWAS analysis for SSE of 2 different splice-sites that identify the same SNP as one of the associations in GWAS. The allelic effects on all both sites are shown. A mutation at the splice acceptor site (At5g65050_28955653) also impacts the downstream splice-donor site c) The SSE differences associated with the associated genotype can be experimentally verified. Common allele is shown in purple and the alternative allele is shown in green. Additional associations detected in the MAF region are given in Supplementary Table 1.

**Supplementary Figure S4.**
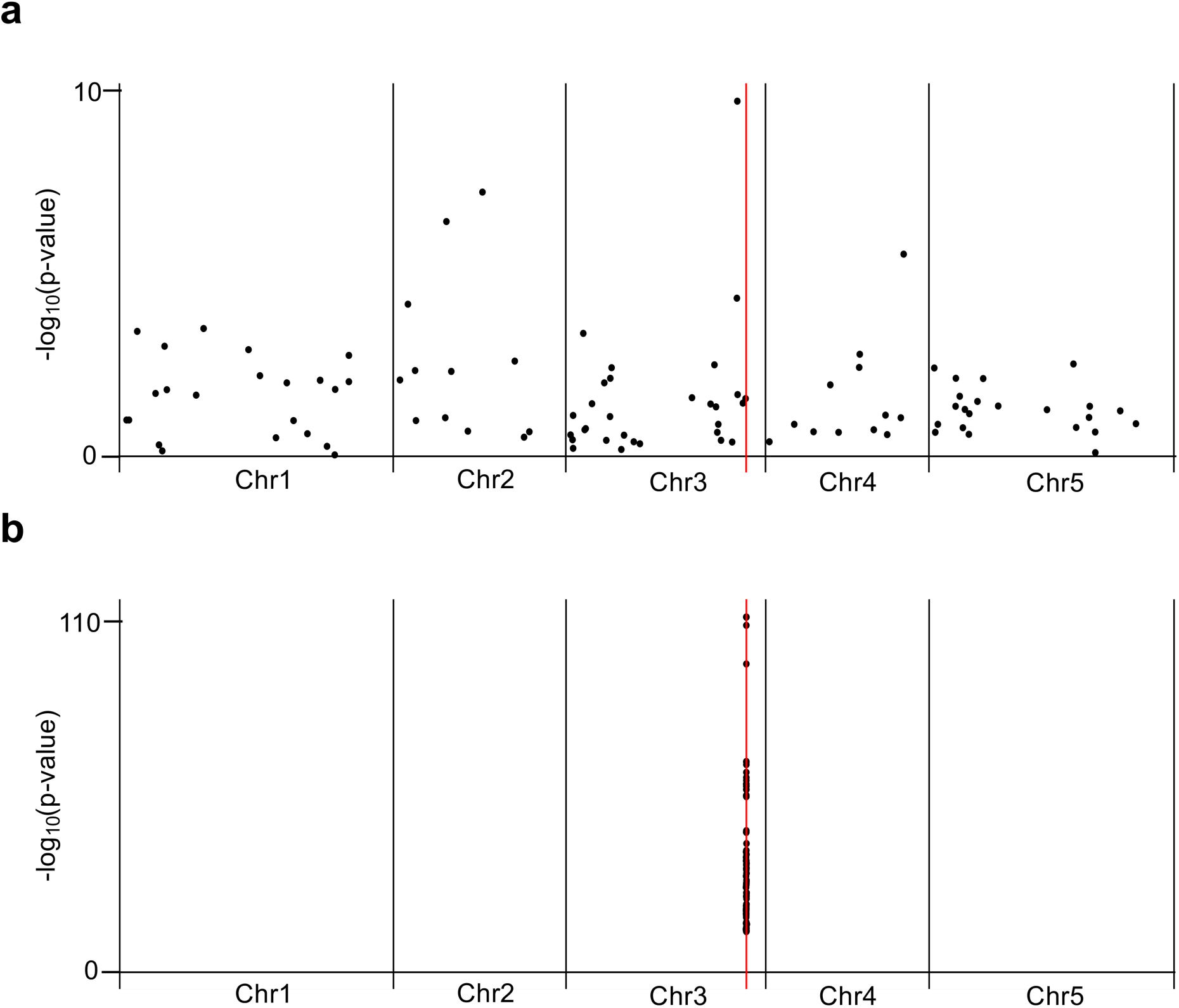
Comparison of associations detected through isoform-based approach and SpliSE-QTL approach. a) Associations reported by Khokhar et al for At3G62190. Only the Top 100 associated SNPs are shown for ease. b) Top 100 associated SNPs through SpliSE QTL approach.

**Supplementary Figure S5.**
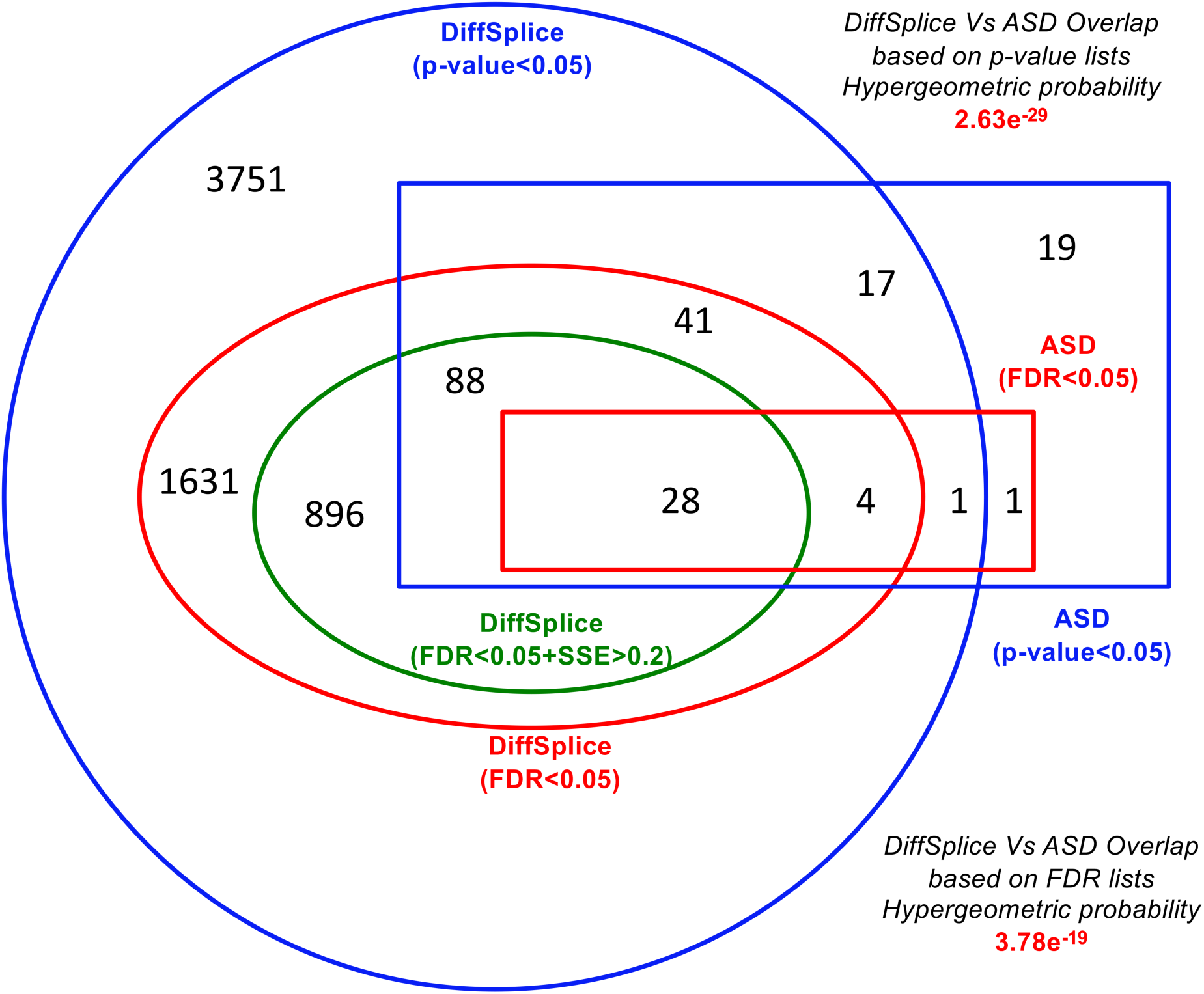
Overlap of the number of genes identified through diffSpliSE analysis with Yan et al study. Ovals represent the data from diffSpliSE and rectangles represent the data from Yan et al. Colours represent equivalent comparisons.

**Supplementary Figure S6.**
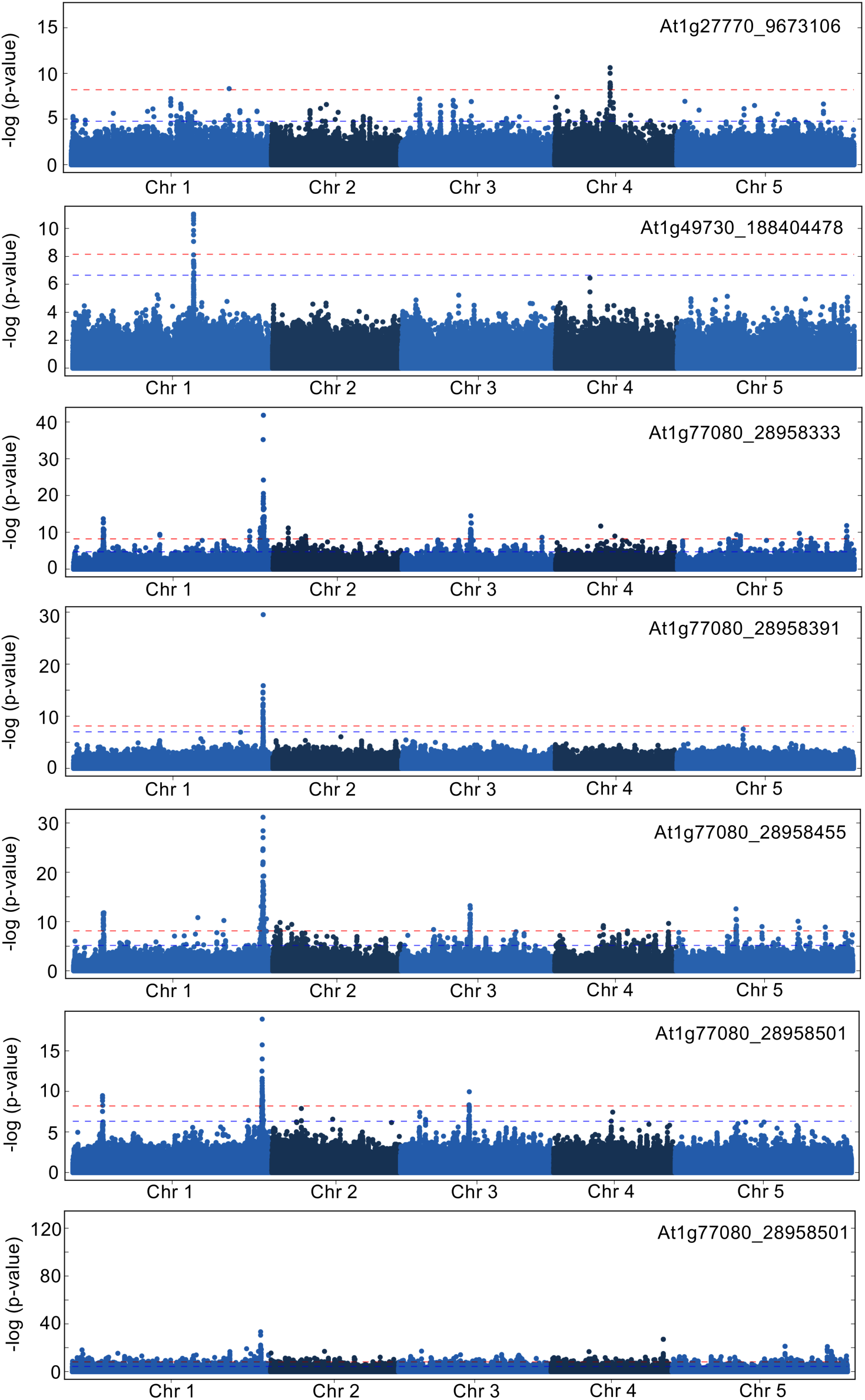
Manhattan plots for detected associations.

**Supplementary Figure S7.**
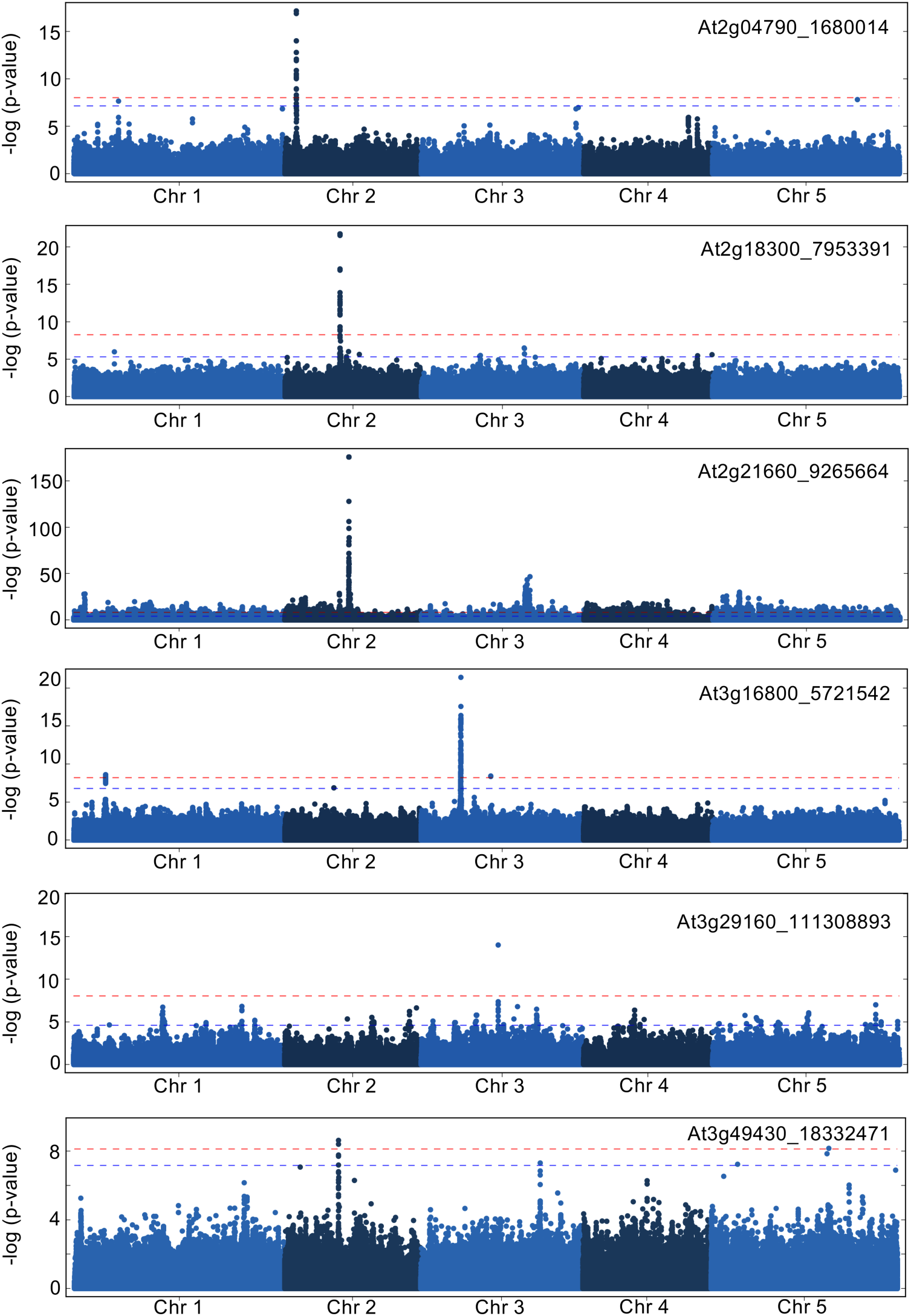
Manhattan plots for detected associations.

**Supplementary Figure S8.**
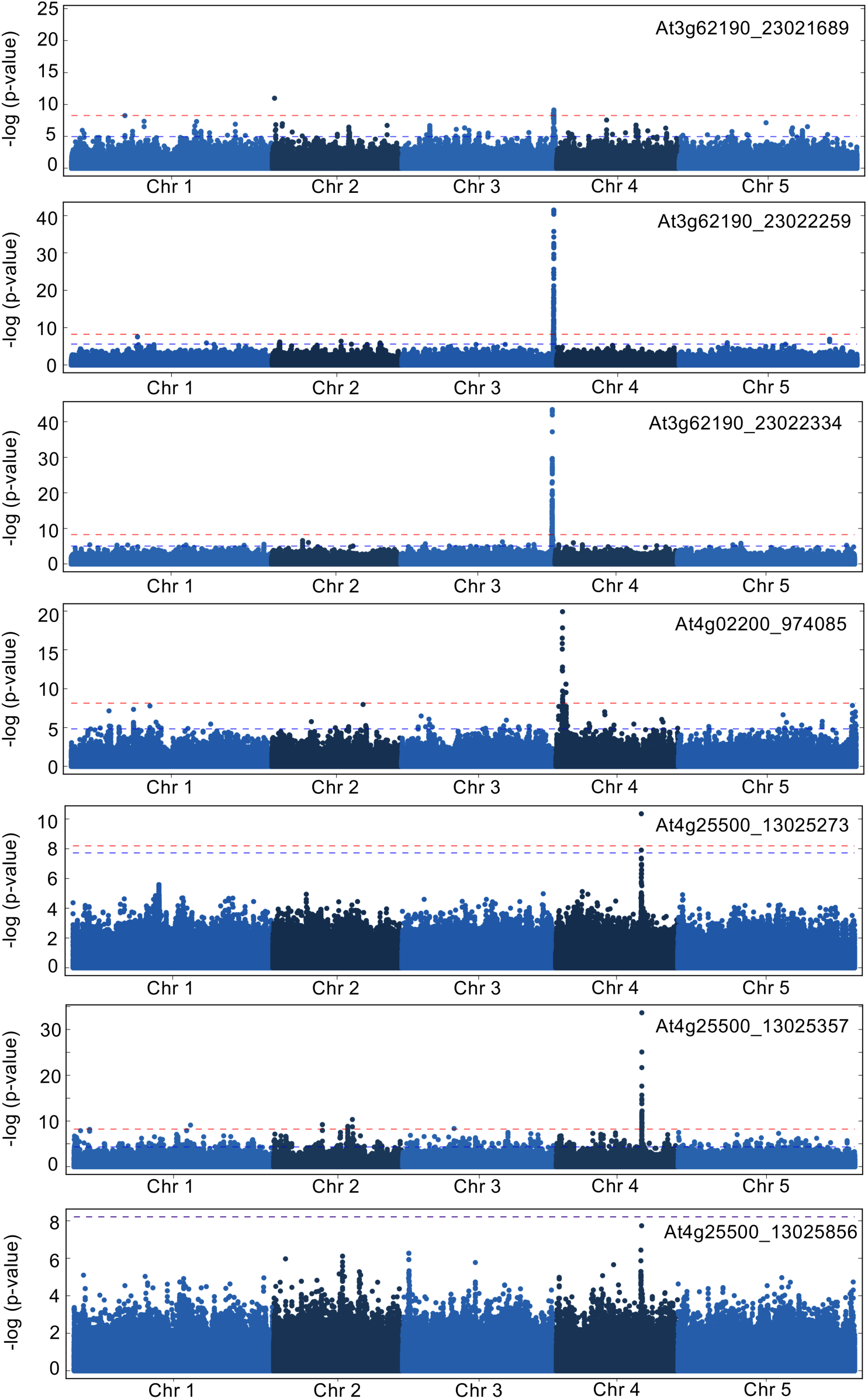
Manhattan plots for detected associations.

**Supplementary Figure S9.**
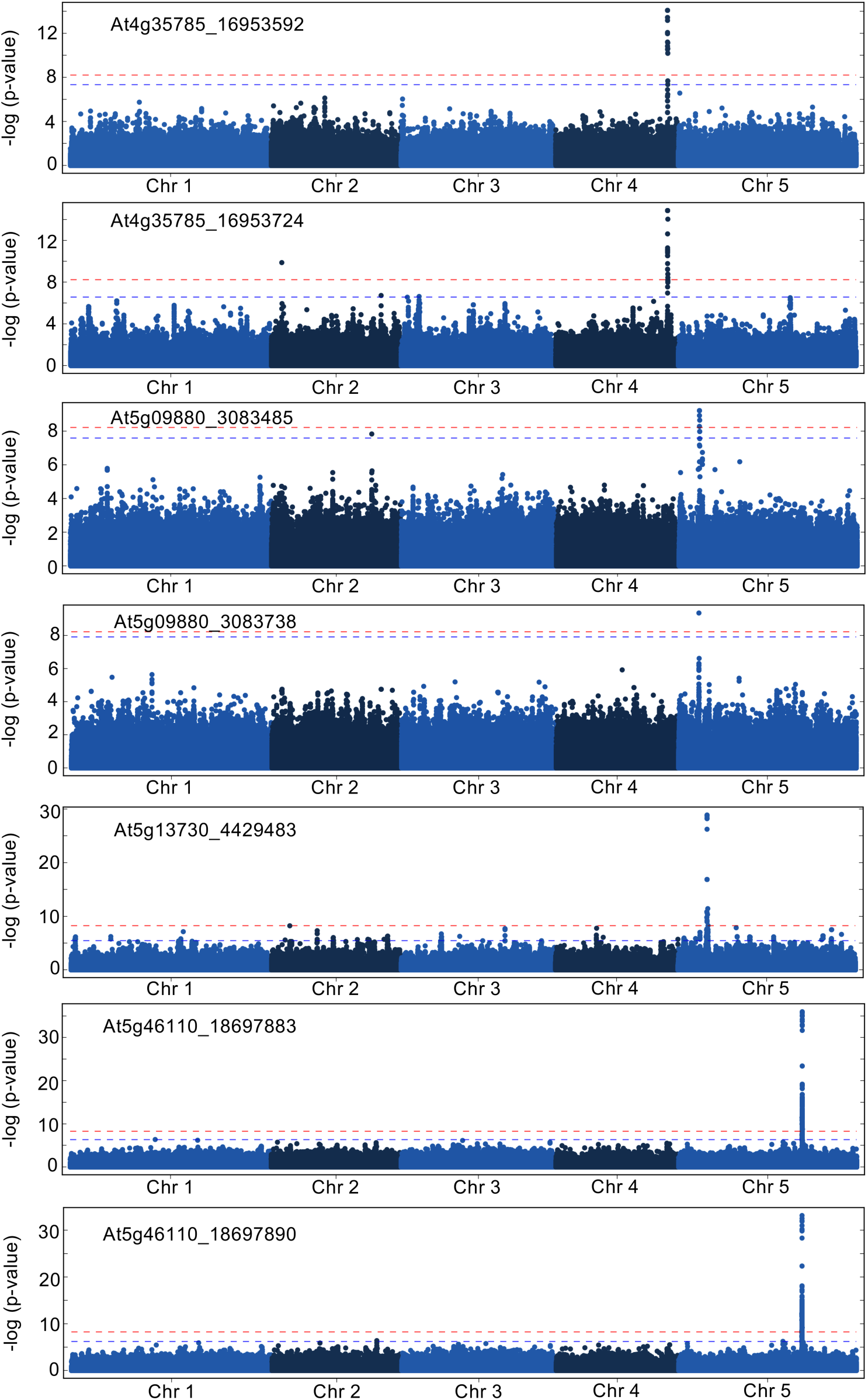
Manhattan plots for detected associations.

**Supplementary Figure S10.**
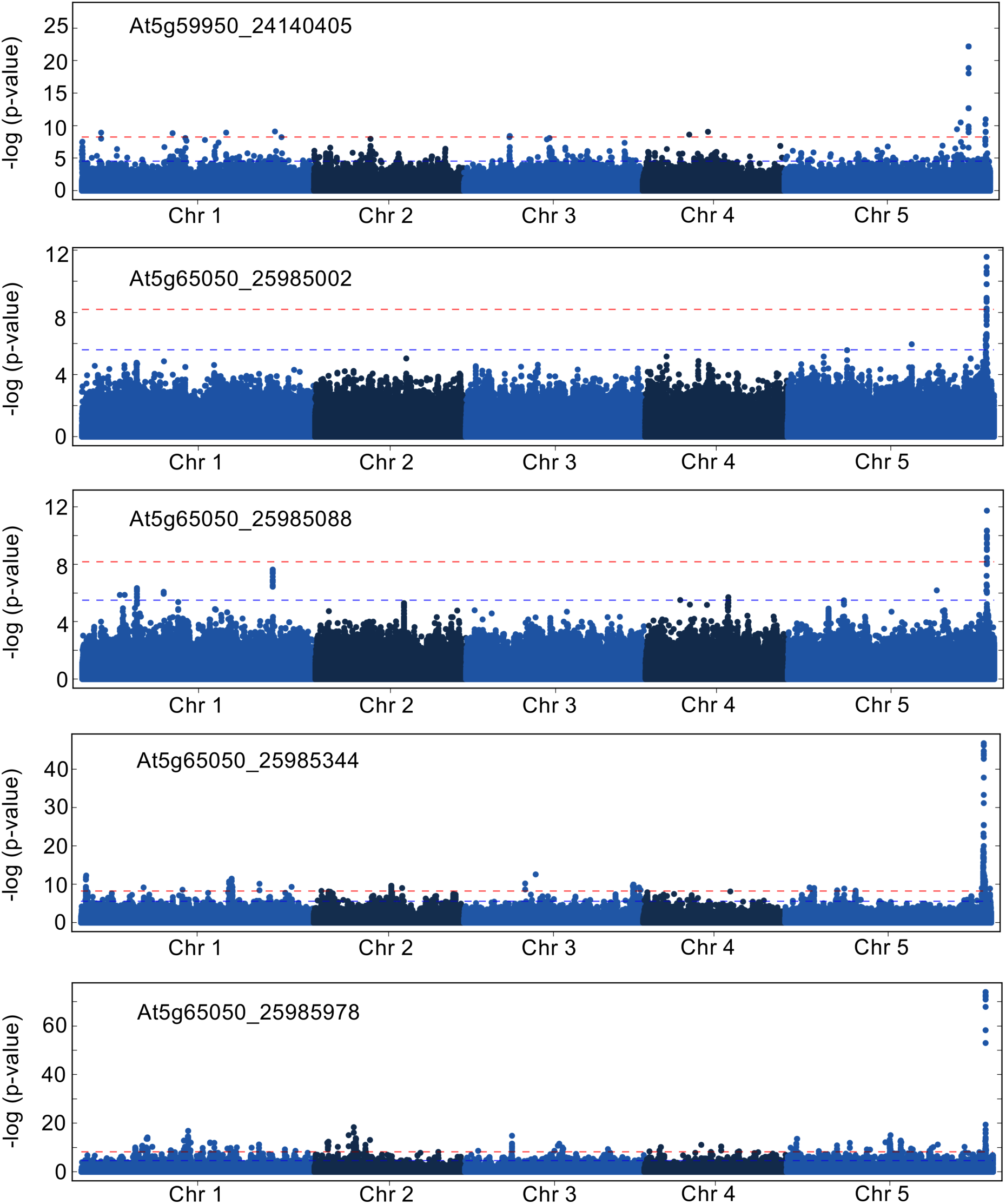
Manhattan plots for detected associations.

**Supplementary Figure S11.**
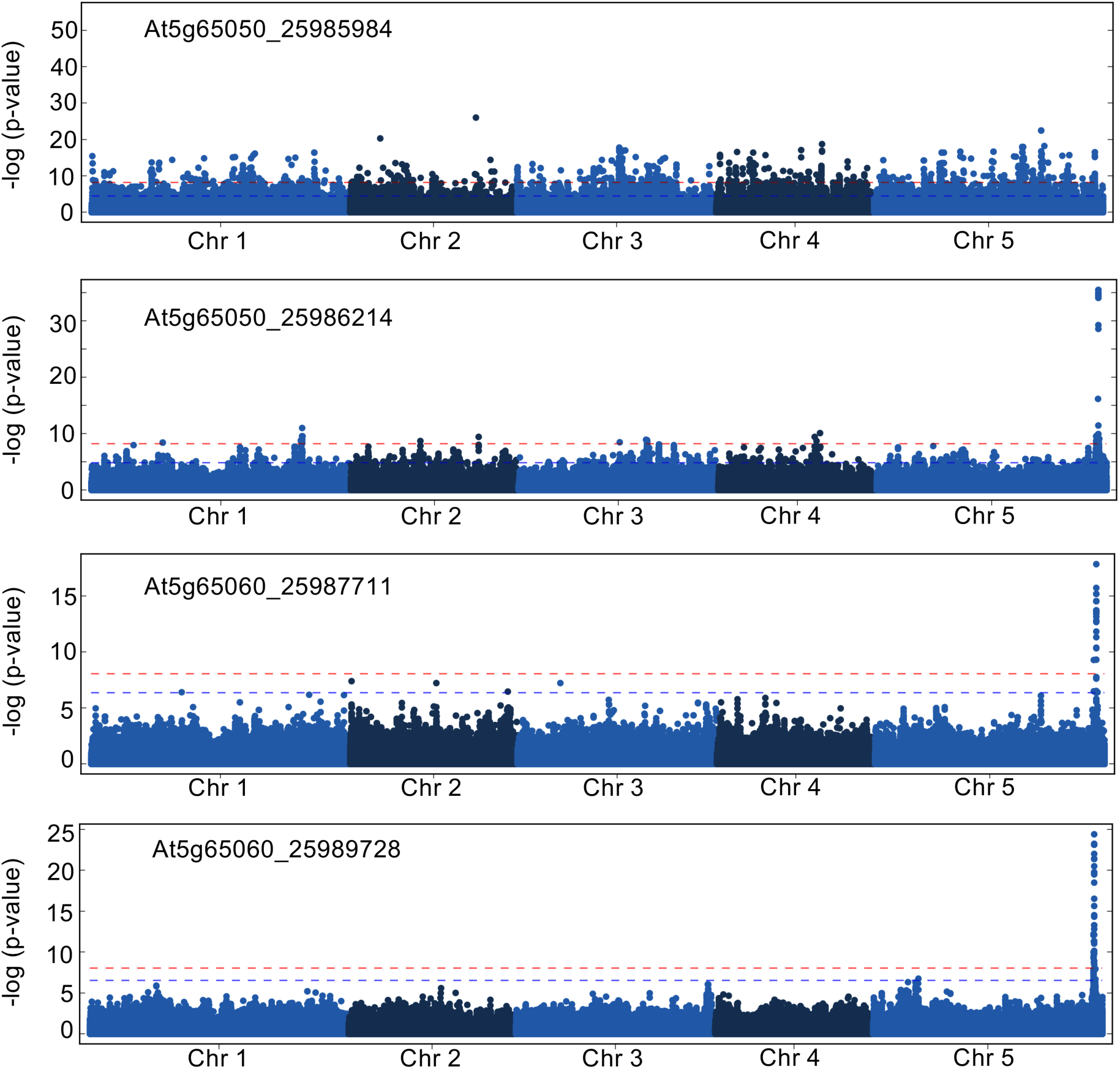
Manhattan plots for detected associations.

### Supplementary Tables.

**Supplementary Table 1.** Summary of the detected associations in SpliSE*-*QTL analysis. When multiple SNPs are in LD, the highest associated SNP refers to the SNP closest to the splice-site among those that display the highest association. In some cases the minor allele count was reduced to capture the SNP that displayed highest association (e.g., AT2G18300_7953316)

**Supplementary Table 2.** Summary of the eQTL analysis on AS-NMD genes

**Supplementary Table 3.** Splice-site strength, heritability, variance and GWAS detection on all detected splice-sites across the *FLM* locus.

**Supplementary Table 4.** Comparison the of GWAS results between isoform-based approach (Khokhar et al, 2019) and our approach, for the same genes.

**Supplementary Table 5.** Primers used for analysis in this study.

## References

1. Li, Y.I. et al. RNA splicing is a primary link between genetic variation and disease. Science 352, 600–4 (2016).

2. Bush, S.J., Chen, L., Tovar-Corona, J.M. & Urrutia, A.O. Alternative splicing and the evolution of phenotypic novelty. Philos Trans R Soc Lond B Biol Sci 372, 20150474 (2017).

3. Laloum, T., Martin, G. & Duque, P. Alternative Splicing Control of Abiotic Stress Responses. Trends Plant Sci 23, 140–150 (2018).

4. Sterne-Weiler, T., Weatheritt, R.J., Best, A.J., Ha, K.C.H. & Blencowe, B.J. Efficient and Accurate Quantitative Profiling of Alternative Splicing Patterns of Any Complexity on a Laptop. Mol Cell 72, 187–200 (2018).

5. Song, Q.A., Catlin, N.S., Brad Barbazuk, W. & Li, S. Computational analysis of alternative splicing in plant genomes. Gene 685, 186–195 (2019).

6. Kalsotra, A. & Cooper, T.A. Functional consequences of developmentally regulated alternative splicing. Nat Rev Genet 12, 715–29 (2011).

7. Szakonyi, D. & Duque, P. Alternative Splicing as a Regulator of Early Plant Development. Front Plant Sci 9, 1174 (2018).

8. Reddy, A.S., Marquez, Y., Kalyna, M. & Barta, A. Complexity of the alternative splicing landscape in plants. Plant Cell 25, 3657–83 (2013).

9. Matera, A.G. & Wang, Z. A day in the life of the spliceosome. Nat Rev Mol Cell Biol 15, 108–21 (2014).

10. Naftelberg, S., Schor, I.E., Ast, G. & Kornblihtt, A.R. Regulation of alternative splicing through coupling with transcription and chromatin structure. Annu Rev Biochem 84, 165–98 (2015).

11. Herzel, L., Ottoz, D.S.M., Alpert, T. & Neugebauer, K.M. Splicing and transcription touch base: co-transcriptional spliceosome assembly and function. Nat Rev Mol Cell Biol 18, 637–650 (2017).

12. Sibley, C.R., Blazquez, L. & Ule, J. Lessons from non-canonical splicing. Nat Rev Genet 17, 407–421 (2016).

13. Baralle, F.E. & Giudice, J. Alternative splicing as a regulator of development and tissue identity. Nat Rev Mol Cell Biol 18, 437–451 (2017).

14. Xu, X. et al. ASF/SF2-regulated CaMKIIdelta alternative splicing temporally reprograms excitation-contraction coupling in cardiac muscle. Cell 120, 59–72 (2005).

15. Salz, H.K. Sex determination in insects: a binary decision based on alternative splicing. Curr Opin Genet Dev 21, 395–400 (2011).

16. Alamancos, G.P., Agirre, E. & Eyras, E. Methods to study splicing from high-throughput RNA sequencing data. Methods Mol Biol 1126, 357–97 (2014).

17. Liu, R., Loraine, A.E. & Dickerson, J.A. Comparisons of computational methods for differential alternative splicing detection using RNA-seq in plant systems. BMC Bioinformatics 15, 364 (2014).

18. Zhang, R. et al. A high quality Arabidopsis transcriptome for accurate transcript-level analysis of alternative splicing. Nucleic Acids Res 45, 5061–5073 (2017).

19. Hooper, J.E. A survey of software for genome-wide discovery of differential splicing in RNA-Seq data. Hum Genomics 8, 3 (2014).

20. Pose, D. et al. Temperature-dependent regulation of flowering by antagonistic *FLM* variants. Nature 503, 414–417 (2013).

21. Sureshkumar, S., Dent, C., Seleznev, A., Tasset, C. & Balasubramanian, S. Nonsense-mediated mRNA decay modulates *FLM*-dependent thermosensory flowering response in Arabidopsis. Nat Plants 2, 16055 (2016).

22. Andreadis, A., Gallego, M.E. & Nadal-Ginard, B. Generation of protein isoform diversity by alternative splicing: mechanistic and biological implications. Annu Rev Cell Biol 3, 207–242 (1987).

23. Breitbart, R.E., Andreadis, A. & Nadal-Ginard, B. Alternative splicing: a ubiquitous mechanism for the generation of multiple protein isoforms from single genes. Annu Rev Biochem 56, 467–495 (1987).

24. Shen, S. et al. rMATS: robust and flexible detection of differential alternative splicing from replicate RNA-Seq data. Proc Natl Acad Sci U S A 111, E5593–601 (2014).

25. Yan, Q., Xia, X., Sun, Z. & Fang, Y. Depletion of Arabidopsis SC35 and SC35-like serine/arginine-rich proteins affects the transcription and splicing of a subset of genes. PLoS Genet 13, e1006663 (2017).

26. Trincado, J.L. et al. SUPPA2: fast, accurate, and uncertainty-aware differential splicing analysis across multiple conditions. Genome Biol 19, 40 (2018).

27. Vaquero-Garcia, J. et al. A new view of transcriptome complexity and regulation through the lens of local splicing variations. Elife 5, e11752 (2016).

28. Li, Y.I. et al. Annotation-free quantification of RNA splicing using LeafCutter. Nat Genet 50, 151–158 (2018).

29. Drechsel, G. et al. Nonsense-mediated decay of alternative precursor mRNA splicing variants is a major determinant of the Arabidopsis steady state transcriptome. Plant Cell 25, 3726–3742 (2013).

30. Kalyna, M. et al. Alternative splicing and nonsense-mediated decay modulate expression of important regulatory genes in Arabidopsis. Nucleic Acids Res 40, 2454–2469 (2012).

31. Reed, R. & Maniatis, T. A role for exon sequences and splice-site proximity in splice-site selection. Cell 46, 681–690 (1986).

32. Eperon, L.P., Estibeiro, J.P. & Eperon, I.C. The role of nucleotide sequences in splice site selection in eukaryotic pre-messenger RNA. Nature 324, 280–282 (1986).

33. Scortecci, K.C., Michaels, S.D. & Amasino, R.M. Identification of a MADS-box gene, *FLOWERING LOCUS M*, that represses flowering. Plant J 26, 229–236 (2001).

34. Kawakatsu, T. et al. Epigenomic Diversity in a Global Collection of *Arabidopsis thaliana* Accessions. Cell 166, 492–505 (2016).

35. Khokhar, W. et al. Genome-Wide Identification of Splicing Quantitative Trait Loci (sQTLs) in Diverse Ecotypes of *Arabidopsis thaliana*. Front Plant Sci 10, 1160 (2019).

36. Monlong, J., Calvo, M., Ferreira, P.G. & Guigo, R. Identification of genetic variants associated with alternative splicing using sQTLseekeR. Nat Commun 5, 4698 (2014).

37. Yang, Q., Hu, Y., Li, J. & Zhang, X. ulfasQTL: an ultra-fast method of composite splicing QTL analysis. BMC Genomics 18, 963 (2017).

38. Wang, X. et al. Cis-regulated alternative splicing divergence and its potential contribution to environmental responses in Arabidopsis. Plant J 97, 555–570 (2019).

39. Zhu, W. et al. Natural Variation Identifies *ICARUS1*, a Universal Gene Required for Cell Proliferation and Growth at High Temperatures in *Arabidopsis thaliana*. PLoS Genet 11, e1005085 (2015).

40. Feng, Y.-Y. et al. RegTools:Integrated analysis of genomic and transcriptomic data for discovery of splicing variants in cancer. BioRxives https://www.biorxiv.org/content/10.1101/436634v2 (2018).

41. Li, H. et al. The Sequence Alignment/Map format and SAMtools. Bioinformatics 25, 2078–2079 (2009).

42. Kim, D. et al. TopHat2: accurate alignment of transcriptomes in the presence of insertions, deletions and gene fusions. Genome Biol 14, R36 (2013).

43. Grimm, D.G. et al. easyGWAS: A Cloud-Based Platform for Comparing the Results of Genome-Wide Association Studies. Plant Cell 29, 5–19 (2017).

44. Seren, U. et al. GWAPP: a web application for genome-wide association mapping in Arabidopsis. Plant Cell 24, 4793–4805 (2012).

45. Robinson, M.D., McCarthy, D.J. & Smyth, G.K. edgeR: a Bioconductor package for differential expression analysis of digital gene expression data. Bioinformatics 26, 139–140 (2010).

